# Rapid and flexible assessment of gene functions in plant cells with particle bombardment and linear DNA

**DOI:** 10.64898/2026.05.17.725698

**Authors:** Pabasara Rasadaree Weerasinghe, Daisuke Tsugama

## Abstract

Biolistic transformation is a versatile tool in plant science, yet high equipment costs and tissue damage from high-pressure gas remain significant barriers. Building on our previously developed “TSGMAC”, a low-cost, helium-free biolistic system, we report three major advancements to enhance its throughput, delivery quality, and quantitative capability. First, a “guide barrel” assembled from commercial DIY fittings was developed; it effectively eliminates physical tissue damage and ensures uniform particle distribution, even in soft tissues like bok choy (*Brassica rapa* subsp. *chinensis*). Second, a rapid gene expression platform using PCR products was characterized. Results demonstrate that linear DNA constructs are efficiently circularized via non-homologous end joining (NHEJ) in plant cells, and protein expression is robust regardless of the relative positions of the promoter, coding sequence, and terminator. This system bypasses time-consuming cloning. Third, a cost-effective, highly sensitive dual-luciferase assay system utilizing teal Luc (teLuc) and inexpensive firefly luciferase (FLuc) inhibitors was established. This integrated workflow enables rapid, quantitative molecular biology using supermarket-obtained materials and standard PCR reagents. Our findings provide a practical foundation for plant scientists, synergistically accelerating gene functional analysis and genetic tool development.

## Introduction

Bolistics is a useful tool in plant science due to their broad host range and procedural simplicity. However, commercial biolistic devices require significant investment costs and the use of high-pressure helium gas, which create barriers to their adoption and maintenance. We previously reported “TSGMAC”, a low-cost, helium-free biolistic system that can be constructed using only a commercial air compressor and DIY parts (Tsugama 2026). The handheld nature of TSGMAC and its simplified particle firing mechanism make it easy to handle, which is a major advantage. To maximize these benefits, effort has been made to diversify the experimental applications and to increase the throughput of the TSGMAC system. During this process, a challenge was encountered when targeting soft tissues, such as bok choy purchased from supermarkets: localized high-pressure airflow caused physical damage to the tissue. Even when the tissue remained intact, the transformed cells were distributed in a characteristic ring-like pattern. In this study, inspired by reports that the gene transfer efficiency of conventional biolistic devices is significantly improved by the use of a “guide barrel” (Thorpe et al. 2025), we report that a “guide barrel” repurposed from a commercial socket reduces physical damage to the target and improves the uniformity of delivery. Furthermore, while the time required for cloning is a bottleneck that hinders research acceleration, we report a gene expression system utilizing PCR products in plants. This system was inspired by expression platforms in budding yeast (*Kluyveromyces marxianus*) and animal cells that utilize PCR products and non-homologous end joining (NHEJ) (Hoshida et al. 2014; Nakamura et al. 2015). Additionally, we report a low-cost and highly sensitive dual-luciferase assay system based on Firefly luciferase (FLuc), an FLuc inhibitor (Bakhtiarova et al. 2006; Auld et al. 2009), and teal Luc (teLuc) (Yeh et al. 2017), developed to leverage these advancements. The establishment of this system enables the rapid execution of quantitative molecular biology experiments using only commercially available plant materials and general-purpose PCR reagents, without the need for expensive or complex capital investment. This can provide a technical foundation that supports scientists conducting research in resource-limited environments.

## Materials and Methods

### Generation of constructs

pGL3-Basic (Promega, Madison, WI, USA) was digested with *Nhe*I and *Xba*I, and the resulting fragment containing the FLuc coding sequence (CDS) was gel-purified using the FastGene Gel/PCR Extraction Kit (Nippon Genetics, Tokyo, Japan). This fragment was inserted into the *Xba*I site of pBS-35S-nYFP-2 (Tsugama et al. 2012a) using the DNA Ligation Kit (Takara Bio, Kusatsu, Japan), generating pBS-35S-FLuc.

pBI121-35SMCS-GFP (Tsugama et al. 2012b) was digested with *Hind*III and *Sal*I, and the fragment containing the cauliflower mosaic virus (CaMV) 35S promoter (P35S) was gel-purified. This fragment was ligated into the *Hind*III-*Sal*I sites of pRL-null (Promega) to generate pRL-35S. A sequence comprising the ampicillin resistance gene (*Amp*^*R*^), a replication origin, the P35S, and the *Renilla* luciferase (RLuc) CDS was amplified by PCR using KOD FX Neo DNA polymerase (Toyobo, Osaka, Japan) with pRL-35S as a template and the primers listed in Supplementary Table S1. Another fragment containing the CaMV 35S terminator (T35S), the bean yellow dwarf virus (BeYDV) Rep/RepA replicase CDS, and the BeYDV long intergenic region (LIR) was amplified using pRN120 (Addgene plasmid #160696; a gift from Daniel Voytas; Maher et al. 2019) as a template. These PCR products were assembled using the In-Fusion Snap Assembly Master Mix (Takara Bio), generating pRL-PT35S-RepA-LIR.

Similarly, a fragment containing the *Amp*^*R*^, replication origin, and P35S was amplified using pRL-35S as a template. The T35S-Rep/RepA-LIR sequence was amplified as described above. The teLuc CDS was amplified using pcDNA3-Antares2 c-Myc (Addgene plasmid #100027; a gift from Huiwang Ai; Yeh et al. 2017) as a template. These fragments were assembled to generate pTeL-PT35S-RepA-LIR.Linear constructs were generated by either standard or overlap PCR using the templates and primers listed in Supplementary Table S2. The preparation workflow, including PCR conditions and polyethylene glycol (PEG)-mediated purification, is detailed in Supplementary Fig. S1.

### GFP and mCherry assays using onion

To evaluate GFP signals from linear constructs, these constructs or pBS-35SMCS-GFP (Tsugama et al. 2012c) were co-bombarded with pBS-35SMCS-mCherry (Tsugama et al. 2013) into onion (*Allium cepa*) epidermal cells using the TSGMAC system as previously described (Tsugama 2026) with minor modifications. Briefly, 250 ng of each construct was mixed with 7.5 μL of gold particles (0.6-μm diameter, 60 mg/mL; InBio Gold, Hurstbridge, Australia) and an equal volume of PEG solution (50% [w/v] PEG3350, 20 mM MgCl_2_). The suspension was incubated for 5 min at room temperature and centrifuged at 10,000 × *g* for 5 s. The gold particles were resuspended in 15 μL of 70% (v/v) ethanol and sonicated for 10 s. Three 5-μL shots were delivered to the epidermal cells of an onion scale leaf. Following bombardment, samples were incubated at 28°C for 16 h.

Fluorescence images were captured using a BX51 microscope equipped with a DP73 CCD camera and cellSens software (Olympus, Tokyo, Japan). GFP images were acquired with a 1-s exposure without a neutral density (ND) filter, while mCherry images were acquired with a 1-s exposure using a U-25ND25 filter. For linear constructs featuring A_19_ or A_50_ at the 3’ ends, experiments were also performed using 750 ng of DNA.

For quantification, apparently intact cells with clear mCherry signals were selected. Cell outlines were manually traced using the Freehand Selection tool in ImageJ (National Institute of Health) and refined using the Fit Spline function. The mean fluorescence intensity was measured within these regions of interest (ROIs). Local background correction was performed by subtracting the mean intensity of a 5-pixel-wide margin surrounding each cell ROI. Data were pooled from at least eight cells per construct. To assess mCherry fused with three tandem repeats of the SV40 nuclear localization signal (mCherry-3NLS; Weerasinghe and Tsugama 2026), the constructs were co-introduced with pBS-35SMCS-GFP. Cells with or without cytosolic mCherry signals were manually counted.

### Dual luciferase and GFP assays using bok choy

Bok choy (*Brassica rapa* subsp. *chinensis*) was obtained from a local supermarket. A guide barrel was constructed by assembling one large fitting (NF-1023; Asoh, Osaka, Japan) and two small fittings (NF-1032) (Fig. 1A). A 1.5 cm × 1.5 cm target area was marked (Fig. 1B) for three shots per construct combination, representing one biological replicate. Particle bombardment was performed as described above, using 400 ng for all RLuc and teLuc constructs and 250 ng for other constructs. The guide barrel was manually fixed to the de Laval nozzle of the TSGMAC during delivery.

**Figure 1.**
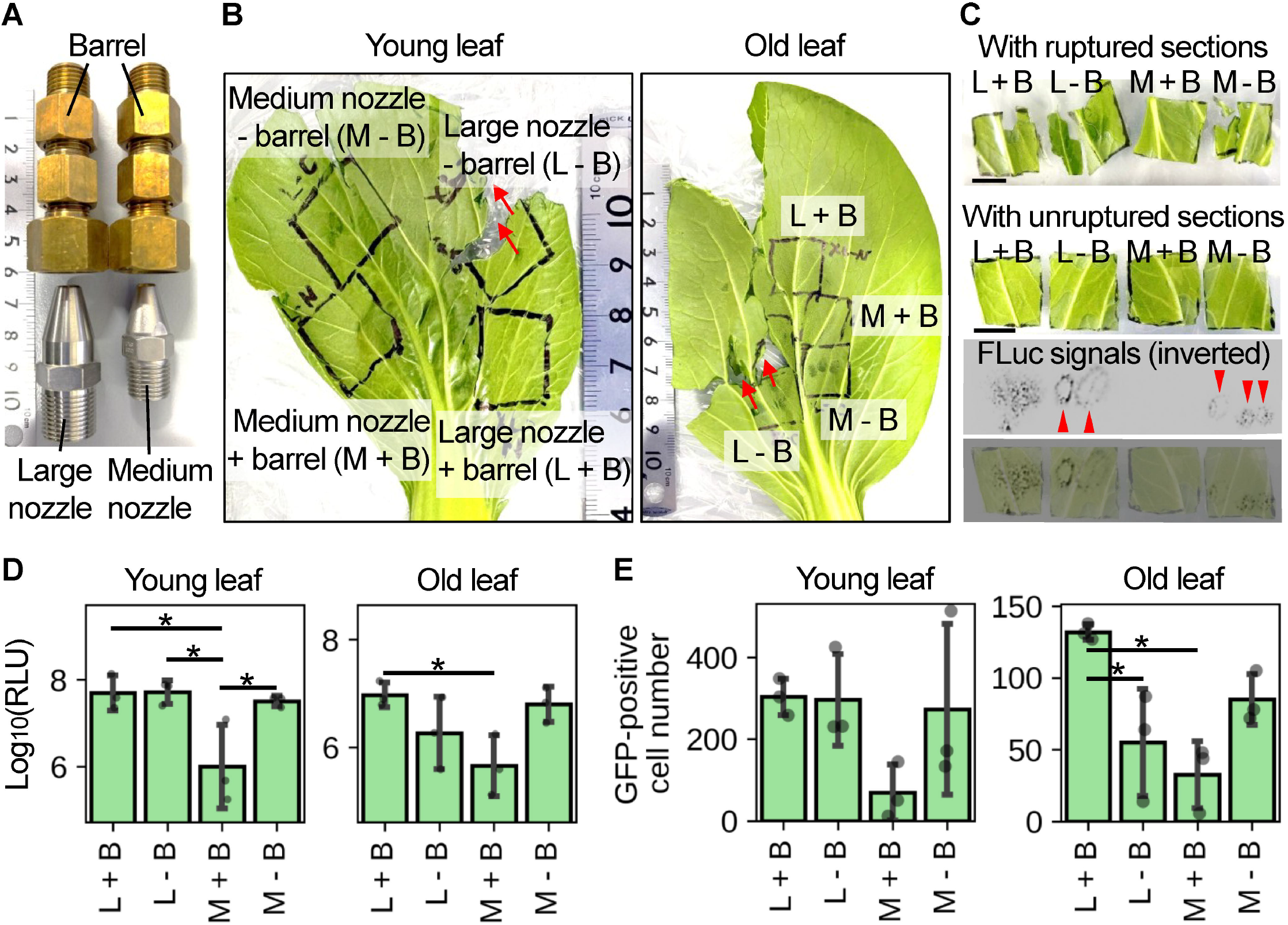
Effect of the guide barrel on tissue damage and transformation patterns. (A) Appearance of the guide barrel (top) and the de Laval nozzles (bottom). (B) Representative images of leaf damage in young (left) and old (right) bok choy leaves bombarded with and without a guide barrel. Conditions are defined as follows: L, large nozzle; M, medium nozzle; B, guide barrel; +, with; –, without. (C) Detection of bioluminescence and fluorescence in leaf sections. The top panel shows a ruptured leaf section; subsequent panels show unruptured sections. Arrowheads indicate ring-like FLuc signal patterns. Scale bars = 1 cm. (D) Quantitative comparison of FLuc signals among different nozzle and barrel combinations. (E) Comparison of the number of GFP-positive cells. In (D) and (E), bars represent mean ± SD with individual data points shown as dot plots. Asterisks indicate significant differences (*P* < 0.05, Tukey’s test).

To evaluate the effect of the guide barrel, the abaxial side of young (inner) and old (outer) leaves were co-transformed with pBS-35S-FLuc and pBS-35SMCS-GFP and incubated for 5 h at room temperature. The target regions were excised using a razor and covered with plastic film to prevent leaf curling. GFP-positive cells were manually counted under a fluorescence microscope. For FLuc activity assays, the Luciferase Assay Reagent (Promega) was diluted 10-fold with distilled water (DW). 125 μL of this solution was applied to each leaf section on a tray, covered with plastic film, and secured with tape (Fig. 1C). Bioluminescence was detected using an ImageQuant LAS 4000 mini imager (GE Healthcare, Chicago, IL, USA) with incremental exposure up to 3 min.

To assess FLuc inhibitors, ataluren (PTC124; Selleck Chemicals, Houston, TX, USA) and resveratrol (Fujifilm Wako Pure Chemical Corp., Osaka, Japan) were used. Ataluren was dissolved in dimethyl sulfoxide (DMSO) (100 μM stock), and resveratrol was dissolved in 99.5% (v/v) ethanol (100 mM stock). Leaves were incubated at room temperature for 16 h post-bombardment. After initial FLuc signal detection, leaf sections were washed in excess DW. Ataluren and resveratrol stocks were diluted 100-fold in Tris-buffered saline (TBS; 150 mM NaCl, 20 mM Tris-HCl, pH 7.6) to final concentrations of 1 μM and 1 mM, respectively. Each leaf section was treated with 200 μL of the diluted inhibitor. Residual FLuc signals were detected as described above. Precipitates were observed in the diluted resveratrol solution, suggesting that the actual concentration was below 1 mM due to partial insolubility.

To compare RLuc- and teLuc-derived signals, 400 ng each of pRL-PT35S-RepA-LIR and pTeL-PT35S-RepA-LIR were introduced into leaf cells as described above. The leaves were incubated for 16 h at room temperature, and sections were prepared as previously detailed. For RLuc detection, the Dual-Glo Stop & Glo Substrate from the Dual-Glo Luciferase Assay System (Promega) was diluted 100-fold with the Dual-Glo Stop & Glo Buffer according to the manufacturer’s instructions. A 125-μL aliquot of the resulting solution was applied to each leaf section. For teLuc detection, the Nano-Glo DLR Stop & Glo Substrate from the Nano-Glo Dual-Luciferase Reporter Assay System (Promega) was similarly diluted 100-fold with the NanoDLR Stop & Glo Buffer, and 125 μL of the solution was applied per leaf section. The resulting bioluminescence was detected as described above.

For dual-luciferase assays, bombarded leaves were incubated, and FLuc activity was inhibited by resveratrol as described above. The resveratrol solution was removed from the leaf sections using paper towels, followed by the application of 125 μL of a teLuc detection solution (100 μM diphenylterazine [DTZ; Selleck Chemicals, prepared from a 10 mM stock in N,N-dimethylformamide] and 1 mM dithiothreitol in TBS) per section. The leaves and solutions were covered with plastic film and subjected to bioluminescence detection as described above.

For signal quantification, bioluminescence images captured with a 3-min exposure for FLuc and a 1-min exposure for teLuc were used. All images within a single experiment were acquired under identical settings to ensure comparability. Using ImageJ, a square region of interest (ROI) approximately 1.5 cm × 1.5 cm (matching the size of the leaf sections) was defined. To determine the local background, the integrated density was first measured in an empty region of the image. The ROI was then moved onto each leaf section to measure its integrated density. The net signal for each leaf section was calculated by subtracting the background integrated density from the total integrated density of the leaf ROI. For FLuc inhibition assays, signals were normalized to the FLuc activity measured before inhibition. For dual-luciferase assays, the net teLuc signal of each leaf section was first normalized to its corresponding FLuc signal to account for variations in transformation efficiency. These FLuc-normalized teLuc activities were then expressed as relative values compared to the activity of the circular construct (pTeL-PT35S-RepA-LIR) control.

## Results and Discussion

### Effects of the guide barrel

The “guide barrel” used in this study was assembled from three commercial fittings (Fig. 1A, top) and was manually pressed against the tip of the de Laval nozzle during delivery. This barrel was tested in combination with medium (M) and large (L) nozzles (Fig. 1A, bottom), which were previously shown to maximize the number of transformed cells in the TSGMAC system (Tsugama 2026). Without the barrel, bok choy leaves, both young (inner) and old (outer), frequently suffered physical damage or tearing; however, the use of the barrel prevented such tissue damage (Fig. 1B).

Regarding the distribution of transformed cells, FLuc-derived signals appeared in a characteristic ring-like pattern when the barrel was absent (Fig. 1C, arrowheads). When the barrel was used, the combination of the medium nozzle and barrel (M + B) yielded almost no detectable signal, whereas the large nozzle and barrel (L + B) produced a uniform distribution of signals across the target area (Fig. 1C). Quantitative analysis of FLuc activity revealed a significant difference between the M + B condition and all other conditions in young leaves, whereas no significant differences were found among the other combinations (Fig. 1D). In the number of GFP-positive cells in old leaves, significant differences were observed between the “L nozzle only (L - B)” and “L + B” conditions, as well as between the “L + B” and “M + B” conditions (Fig. 1E). These results suggest that while the use of a guide barrel in the TSGMAC system does not necessarily increase the total number of transformed cells, it promotes a more uniform delivery pattern and protects soft tissues from high-pressure airflow.

### Characterization of the PCR-product-based gene expression system in plants

To evaluate the impact of the arrangement of the promoter, marker gene coding sequence (CDS), and terminator, as well as the effect of a poly(A) tail, on gene expression efficiency, various PCR-derived linear constructs were introduced into onion epidermal cells (Fig. 2A). Interestingly, constructs with the arrangements “terminator–promoter–GFP” and “GFP–terminator–promoter” showed GFP induction efficiencies comparable to the standard “promoter–GFP–terminator” configuration (Fig. 2B, first three constructs).

**Figure 2.**
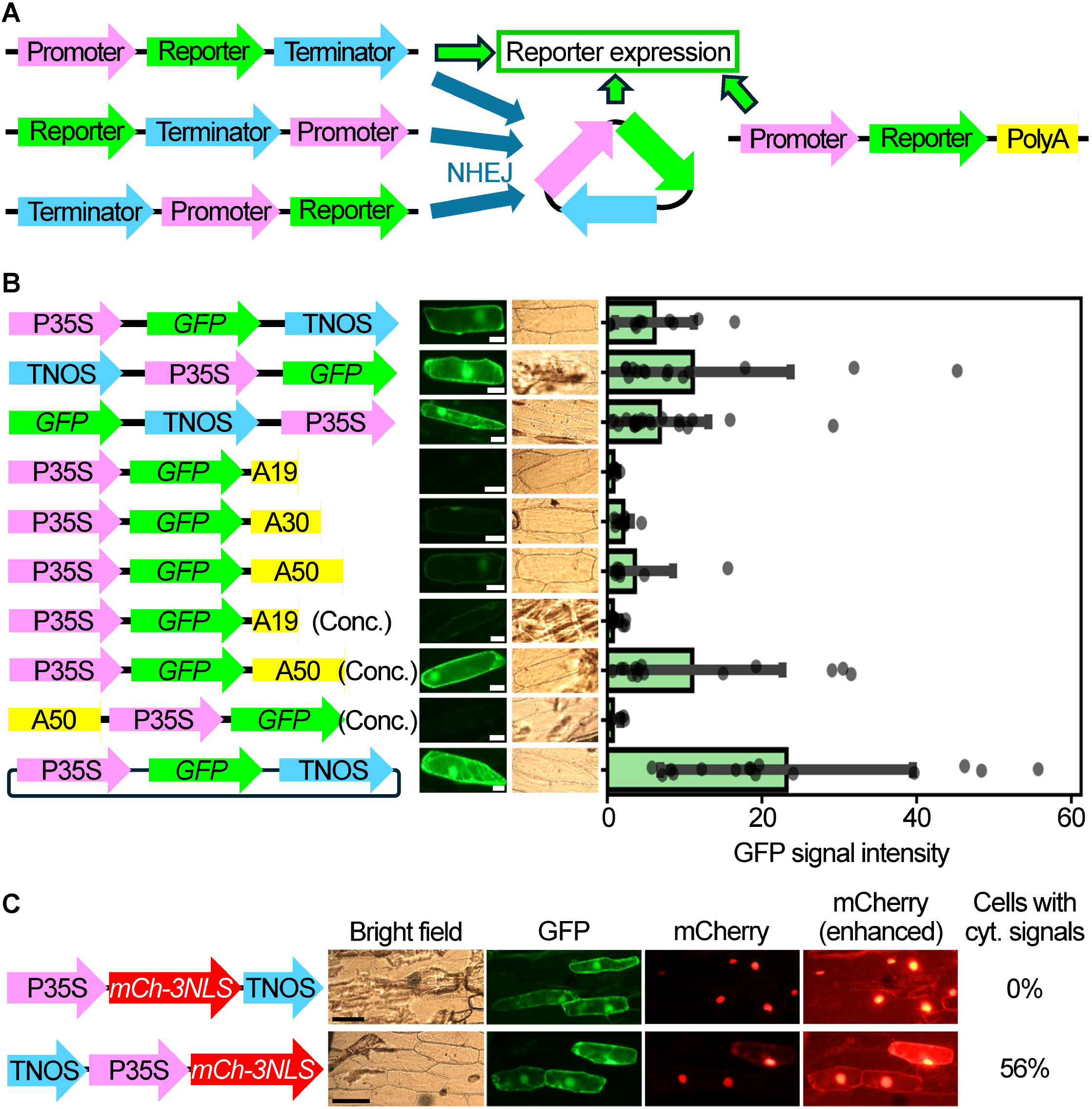
Characterization of the PCR-product-based gene expression system. (A) Conceptual diagrams. Left: circularization of three different linear constructs via NHEJ results in structurally equivalent circular DNA. Right: a linear DNA construct with a 3’ poly A tail. (B) Schematic diagrams of linear constructs (left), representative fluorescence and bright-field images (middle), and quantitative GFP signal analysis (right). Scale bars = 50 μm. Bars for the quantitative data represent mean ± SD with individual data points shown as dot plots. (C) Schematic diagrams of mCherry-3NLS (mCh-3NLS) constructs (left), representative images (middle; bright-field, GFP signals, mCh-3NLS signals, and enhanced mCh-3NLS signals) and the percentage of fluorescent cells with cytosolic (cyt.) mCherry signals (right). The enhanced images were used to detect the cytosolic mCherry signals from the construct with *mCh-3NLS* at the 3’ terminus. Scale bars = 250 μm.

In experiments using “promoter–GFP–poly(A)” constructs, longer poly A tails (A_50_) resulted in higher GFP induction efficiency than shorter ones (A_19_ and A_30_). When using the A_50_ construct, increasing the amount of DNA further enhanced GFP induction. In contrast, almost no GFP signal was detected from the “A_50_–promoter–GFP” arrangement (Fig. 2B, following six constructs). These findings suggest that, similar to observations in *K. marxianus* and animal cells (Hoshida et al. 2014; Nakamura et al. 2015), exogenous linear DNA is efficiently circularized via NHEJ in plant cells. They also highlight the critical role of the terminator and subsequent polyadenylation in achieving protein expression.

To further characterize this process, mCherry-3NLS (mCherry fused with three tandem repeats of the SV40 NLS) was expressed using two linear configurations: “promoter–mCh-3NLS–terminator” and “terminator–promoter–mCh-3NLS.” While mCherry signals from the former were limited to the nucleus, signals from the latter were observed in the cytosol (though weakly) in 56% of the fluorescent cells (Fig. 2C). This suggests that several dozen base pairs at the ends of the linear DNA can be degraded before NHEJ-mediated circularization occurs, potentially compromising the function of at least one of the three NLS repeats.

### Establishment of a biolistic dual-luciferase assay system

A three-step biolistic dual-luciferase assay system (FLuc detection, FLuc inhibition, and teLuc detection) was established using bok choy leaves as the target material (Fig. 3A). Regarding FLuc inhibition, our preliminary experiments using commercial dual-luciferase reagents showed non-negligible residual FLuc signals. Therefore, ataluren and resveratrol, both previously reported to inhibit FLuc activity (reference), were tested. Upon application of either inhibitor, FLuc signals were almost completely abolished, even when measured immediately after treatment (Fig. 3B). While ataluren was dissolved in DMSO, we selected resveratrol (dissolved in ethanol) for routine analysis due to its convenience, as the stock solution remains liquid even under cold storage. For the second luciferase, teLuc was adopted instead of the commonly used RLuc, as teLuc exhibited significantly stronger signals in our system (Fig. 3C).

**Figure 3.**
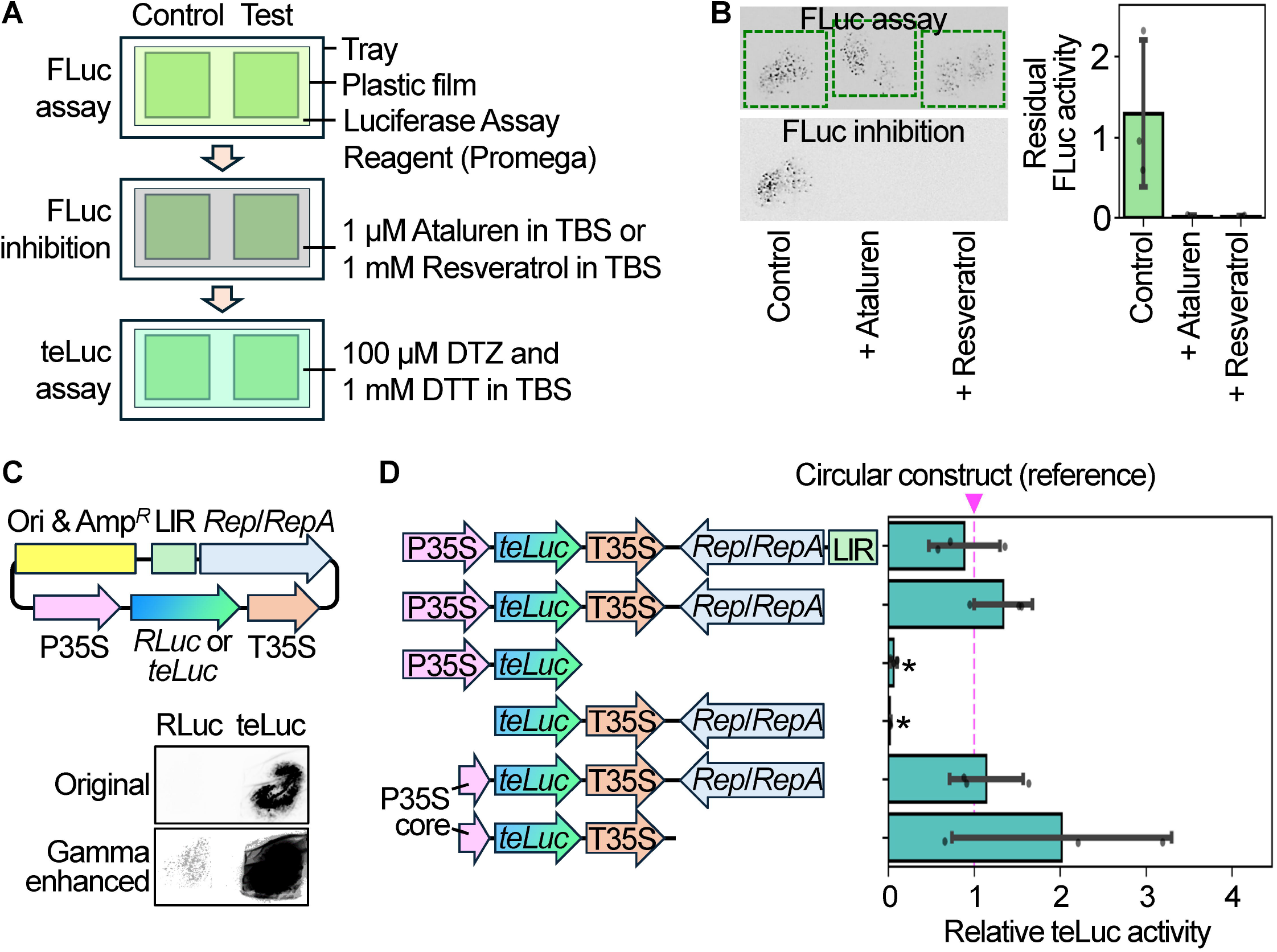
Establishment of a biolistic dual-luciferase assay system. (A) Schematic overview of the three-step assay and the reagents used. (B) Efficacy of FLuc inhibitors. Representative inverted bioluminescence images (left) and quantitative results (right) are shown. In the images, dashed squares (1.5 cm × 1.5 cm) indicate the approximate positions of the leaf sections. (C) Comparison of RLuc and teLuc circular constructs. Schematics (top) and representative inverted bioluminescence images (bottom) are shown. The gamma-enhanced image is presented to show the presence of the RLuc signals. (D) Schematic diagrams of linear teLuc constructs used for the dual-luciferase assay (left) and their quantitative results (right). teLuc activities were normalized to FLuc and expressed relative to the circular construct control. Asterisks indicate significant differences from the reference (circular construct) value of 1.0 (*P* < 0.05, Student’s *t*-test). In (B) and (D), bars represent mean ± SD with individual data points shown as dot plots.

As a practical application, the effects of various regulatory elements on gene expression were evaluated using this assay. For this, a circular plasmid containing a geminivirus replicon (Rep/RepA and the LIR from BeYDV (Maher et al. 2019)) and an expression cassette (P35S–teLuc CDS–T35S), as well as several linear constructs amplified by PCR from this plasmid, was generated. The absence of teLuc signals in constructs lacking either the P35S or T35S reaffirmed the necessity of both the promoter and terminator for protein expression. Unexpectedly, a minimal construct containing only the P35S core promoter, the teLuc CDS, and the T35S induced teLuc expression levels comparable to those of other constructs, including the full circular plasmid (Fig. 3D). Since we standardized the mass of the teLuc constructs for bombardment, smaller constructs were delivered at a higher molar concentration. This higher molecular abundance per cell may have compensated for the lack of upstream enhancer regions or the viral replicon. Regarding the geminivirus replicon, although the LIR is known to function as both an origin of replication and a promoter (Mor et al. 2003), it is possible that it did not function optimally within our specific construct configurations. Furthermore, because signals were detected 16 h post-bombardment, the accumulation of teLuc protein might have reached saturation regardless of promoter strength or DNA copy number. To improve this assay for detecting relatively small differences in promoter activity, the use of protein-destabilizing motifs (e.g., PEST and C9 domains (Yasunaga et al. 2015)) or the optimization of incubation times may be beneficial. Given that the reagents required for this assay are commercially available and sufficient for numerous reactions per purchase, we believe this system offers a cost-effective solution for quantitative molecular biology.

The PCR-product-based direct expression system, which enables efficient gene induction even when the promoter, CDS, and terminator arrangements are altered (particularly with the CDS positioned proximal to the ends), enhances the flexibility of transgene design and reduces the time required for functional evaluation. The introduction of a guide barrel assembled from commercial DIY parts addresses the inherent physical challenges of handheld biolistic devices, namely, tissue damage and delivery non-uniformity, thereby enabling precise analysis using commercially available leafy vegetables. By utilizing relatively inexpensive compounds and teLuc, a cost-effective and sensitive dual-luciferase assay system has been established. These advancements are expected to synergistically accelerate gene functional analysis and the development of novel genetic tools through efficient mutagenesis, domain addition, and domain substitution.

## Acknowledgments

The authors also acknowledge the use of the computational resources provided by The University of Tokyo.

## Author contribution statement

PW and DT conceived the study and designed the experiments. PW and DT performed the experiments and data analysis. Both authors contributed to writing and revising the manuscript.

## Data availability

All data generated or analyzed during this study are included in the published article and its Supplementary files.

## Description of supplementary file

### Supplementary_tables_NHEJ_DL.xlsx

Supplementary Tables S1 and S2, containing lists of primers used for the generation of circular and linear constructs, respectively.

### Supplementary_figure_NHEJ_DL.pdf

Supplementary Fig. S1, illustrating the workflow for linear construct preparation.

## References

Auld DS, Thorne N, Maguire WF, Inglese J (2009) Mechanism of PTC124 activity in cell-based luciferase assays of nonsense codon suppression. Proc Natl Acad Sci U S A 106:3585–3590. 10.1073/pnas.0813345106

Bakhtiarova A, Taslimi P, Elliman SJ, Kosinski PA, Hubbard B, Kavana M, Kemp DM (2006) Resveratrol inhibits firefly luciferase. Biochem Biophys Res Commun 351:481–484. 10.1016/j.bbrc.2006.10.057

Hoshida H, Murakami N, Suzuki A, Tamura R, Asakawa J, Abdel-Banat BM, Nonklang S, Nakamura M, Akada R (2014) Non-homologous end joining-mediated functional marker selection for DNA cloning in the yeast *Kluyveromyces marxianus*. Yeast 31:29–46. 10.1002/yea.2993

Maher MF, Nasti RA, Vollbrecht M, Starker CG, Clark MD, Voytas DF (2019) Plant gene editing through de novo induction of meristems. Nat Biotechnol 38:84–89. 10.1038/s41587-019-0337-2

Mor TS, Moon YS, Palmer KE, Mason HS (2003) Geminivirus vectors for high-level expression of foreign proteins in plant cells. Biotechnol Bioeng 81:430–437. 10.1002/bit.10483

Nakamura M, Suzuki A, Akada J, Tomiyoshi K, Hoshida H, Akada R (2015) End joining-mediated gene expression in mammalian cells using PCR-amplified DNA constructs that contain terminator in front of promoter. Mol Biotechnol 57:1018–1029. 10.1007/s12033-015-9890-1

Thorpe C, Luo W, Ji Q, Eggenberger AL, Chicowski AS, Xu W, Sandhu R, Lee K, Whitham SA, Qi Y, Wang K, Jiang S (2025) Enhancing biolistic plant transformation and genome editing with a flow guiding barrel. Nat Commun 16:5624. 10.1038/s41467-025-60761-x

Tsugama D, Liu H, Liu S, Takano T (2012a) *Arabidopsis* heterotrimeric G protein β subunit interacts with a plasma membrane 2C-type protein phosphatase, PP2C52. Biochim Biophys Acta 1823:2254–2260. 10.1016/j.bbamcr.2012.10.001

Tsugama D, Liu S, Takano T (2012b) Drought-induced activation and rehydration-induced inactivation of MPK6 in Arabidopsis. Biochem Biophys Res Commun 426:626–629. 10.1016/j.bbrc.2012.08.141

Tsugama D, Liu S, Takano T (2012c) A putative myristoylated 2C-type protein phosphatase, PP2C74, interacts with SnRK1 in Arabidopsis. FEBS Lett 586:693–698. 10.1016/j.febslet.2012.02.019

Tsugama D, Liu S, and Takano T (2013) A bZIP protein, VIP1, interacts with Arabidopsis heterotrimeric G protein β subunit, AGB1. Plant Physiol Biochem 71:240–246. 10.1016/j.plaphy.2013.07.024

Tsugama D (2026) A tool to shoot genes with massive air from a compressor (TSGMAC). bioRxiv. 2026.03.24.713841. 10.64898/2026.03.24.71384

Yasunaga M, Murotomi K, Abe H, Yamazaki T, Nishii S, Ohbayashi T, Oshimura M, Noguchi T, Niwa K, Ohmiya Y, Nakajima Y (2015) Highly sensitive luciferase reporter assay using a potent destabilization sequence of calpain 3. J Biotechnol 194:115–123. 10.1016/j.jbiotec.2014.12.004

Yeh HW, Karmach O, Ji A, Carter D, Martins-Green MM, Ai HW (2017) Red-shifted luciferase-luciferin pairs for enhanced bioluminescence imaging. Nat Methods 14:971–974. 10.1038/nmeth.4400

