## Supplementary Fig. S1 for "Rapid and flexible assessment of gene functions in plant cells with particle bombardment and linear DNA"

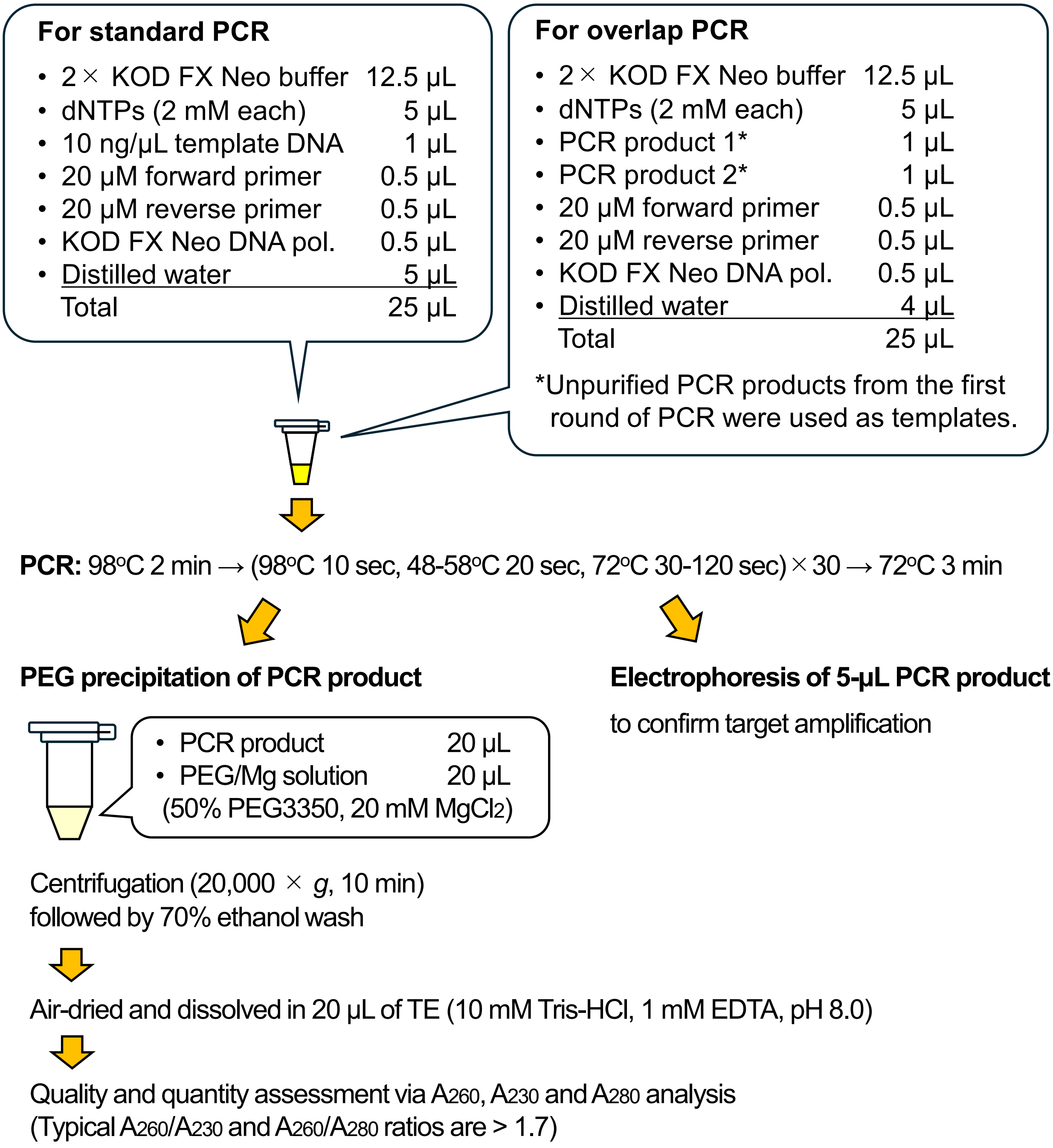


**Supplementary Fig. S1. Flowchart of linear construct preparation via PCR and PEG-mediated purification.**
